# The effect of nutrient availability on global forest carbon balance is uncertain

**DOI:** 10.1101/012476

**Authors:** Enzai Du

## Abstract

To the Editor — Fernández-Martínez *et al.* ^1^ show a chief determinant of nutrient availability on net ecosystem production (NEP) and ecosystem carbon-use efficiency (CUEe, the ratio of NEP to gross primary production i.e. GPP) in global forests. However, their conclusions depend on an improper treatment of differences in the GPP range of nutrient-rich and nutrient-poor forests (uneven sampling effect) and outliers. A statistical re-analysis of their datasets, while *simultaneously* excluding the uneven sampling effect and outliers, indicates no significant control of nutrient availability on carbon (C) balance.

---

First, NEP and CUEe both have a non-linear relationship against GPP (Fig. 1a and 1b) and this indicates that an uneven sampling effect can result in misleading conclusions. Taking nutrient-poor forests as an example, CUEe within the GPP range of 1000∼2000 g C m^-2^ yr^-1^ (16 ± 3%; mean ± s.e.m.) is significantly higher than that within the whole GPP range (6 ± 4%) (*t*-test, p < 0.05). A generalized linear model (GLM) analysis indicates that differences of GPP ranges (e.g. 1000 ∼ 2 000 g C m^-2^ yr^-1^ *vs.* whole GPP range) significantly affect NEP (p < 0.05). Therefore, statistical analysis of Fernández-Martínez *et al.*^1^ should have been based on a same GPP range to exclude the uneven sampling effect.

**Figure 1.**
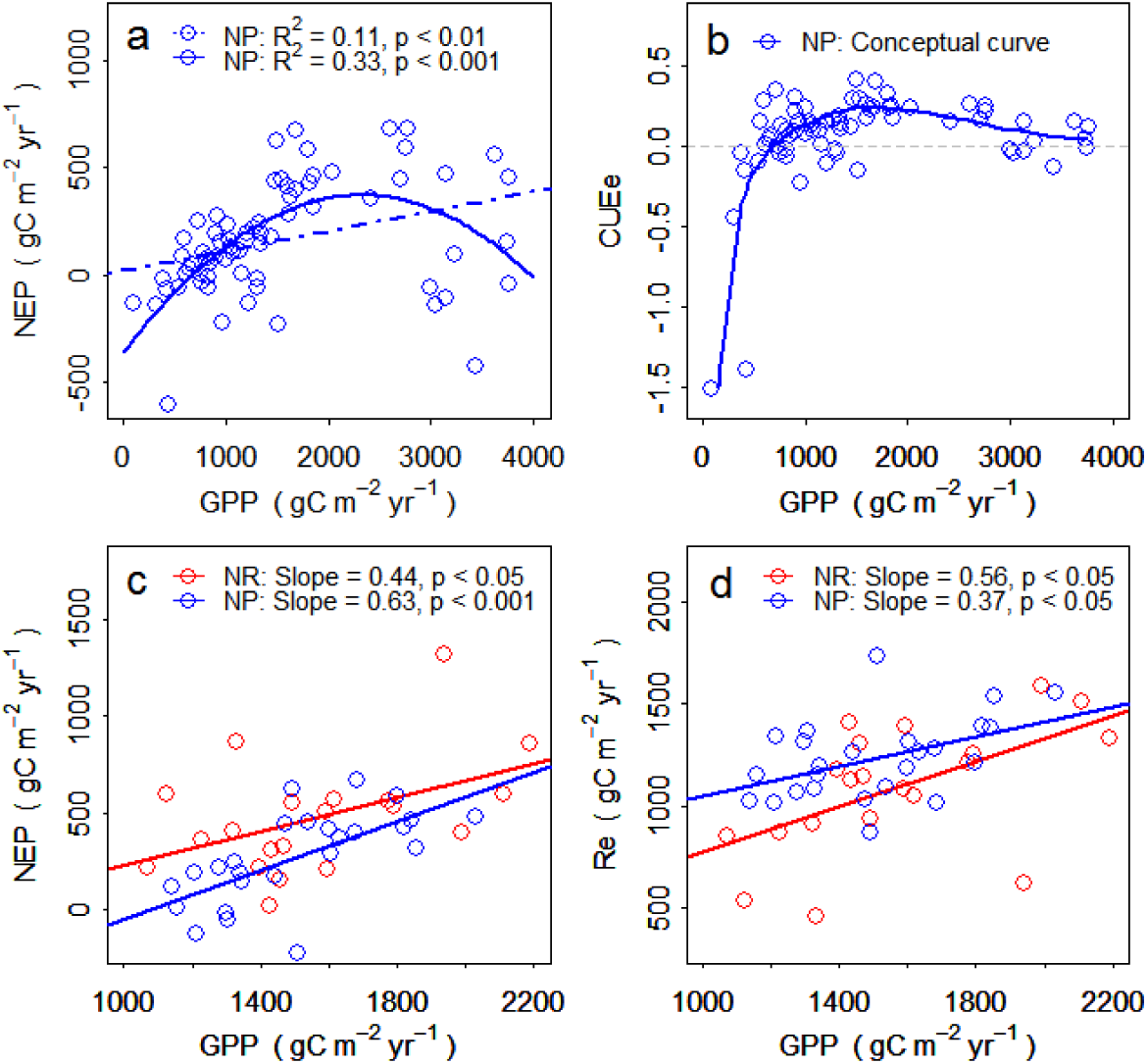
NEP (g C m^-2^ yr^-1^), CUEe, and Re (g C m^-2^ yr^-1^) against GPP (g C m^-2^ yr^-1^) in nutrient-rich (NR) and nutrient-poor (NP) forests. (a) Change in NEP against GPP within whole GPP range in nutrient-poor forests, (b) non-linear conceptual model of CUEe against GPP based on dataset in nutrient-poor forests, (c) comparison of the slope of NEP against GPP, and (d) comparison of the slope of Re against GPP in nutrient-rich and nutrient-poor forests.

Second, three very young forests (< 5 years) with extremely high GPP and NEP are likely outliers, because young forests commonly have low GPP and NEP^2^^,^ ^3^. When excluding these outliers, the slope of NEP against GPP within a common GPP range (1000 ∼ 2200 g C m^-2^ yr^-1^) showed no significant difference (p = 0.49) between nutrient-rich forests (slope = 0.44, p < 0.05) and nutrient-poor forests (slope = 0.63, p < 0.001) (Fig. 1c). The slope of ecosystems respiration (Re) against GPP for nutrient-rich forests (slope = 0.56, p < 0.05) also showed no significant difference (p = 0.85) from that for nutrient-poor forests (slope = 0.37, p < 0.05) (Fig. 1d). These results indicate that nutrient-rich and nutrient-poor forests do not show significant difference in their allocation of GPP to NEP.

Statistical analyses by Fernández-Martínez *et al.* ^1^ have never *simultaneously* excluded the uneven sampling effect and outliers. When doing so, a GLMs analysis indicates that nutrient availability (p = 0.26) and nutrient*GPP interaction (p = 0.49) both exert no significant control on NEP.

Moreover, I propose a non-linear conceptual model of CUEe against GPP (Fig. 1b). Youngest forests commonly show very low GPP and negative CUEe because of higher Re than GPP ^2^, and then CUEe increases rapidly with growing GPP to a critical point which is C neutral. CUEe continues to increase but starts to slow down at a certain stage when nutrient limitation is intensified by biomass nutrient accumulation ^4^, and further it reaches a maximum after which CUEe declines slowly due to increasing allocation of GPP to Re^5^. This conceptual model implies that, instead of nutrient availability, GPP and stand age may jointly determine C allocation of GPP to NEP in global forests.

